# The effects of mutational processes and selection on driver mutations across cancer types

**DOI:** 10.1101/149096

**Authors:** Daniel Temko, Ian PM Tomlinson, Simone Severini, Benjamin Schuster-Böckler, Trevor A Graham

**Affiliations:** Evolution and Cancer laboratory, Barts Cancer Institute, QMUL, Charterhouse Sq, London, EC1M 6BQ.; Department of Computer Science, University College London, London, WC1E 6BT.; Centre for Mathematics and Physics in the Life Sciences and Experimental Biology (CoMPLEX), University College London, London, WC1E 6BT.; Institute of Cancer and Genomic Sciences, University of Birmingham; Institute of Natural Sciences, Shanghai Jiao Tong University.; Ludwig Institute for Cancer Research, University of Oxford, Oxford, OX3 7DQ.

## Abstract

Epidemiological evidence has long associated environmental mutagens with increased cancer risk. However, links between specific mutation-causing processes and the acquisition of individual driver mutations have remained obscure. Here we have used public cancer sequencing data to infer the independent effects of mutation and selection on driver mutation complement. First, we detect associations between a range of mutational processes, including those linked to smoking, ageing, APOBEC and DNA mismatch repair (MMR) and the presence of key driver mutations across cancer types. Second, we quantify differential selection between well-known alternative driver mutations, including differences in selection between distinct mutant residues in the same gene. These results show that while mutational processes play a large role in determining which driver mutations are present in a cancer, the role of selection frequently dominates.

## INTRODUCTION

Environmental mutagens have long been associated with cancer risk^1-3^, but links between mutagens and the generation of specific pathological mutations have remained obscure. A landmark study by Alexandrov et al.^4,5^ identified distinct mutational”signatures”, each representing distinct mutagenic processes, many of which are attributable to environmental mutagens. Each signature consists of the frequency of mutations in 96 “channels” of somatic single nucleotide substitution variants (SNVs) in the contexts of the two flanking bases. The study described 21 different mutational signatures, each characterised by different proportions of the 96 types. Subsequently more than 30 signatures, many with tumour type-specificity, have been reported^6-11^.

The likelihood of acquisition of specific cancer-causing mutations^12^, hereafter referred to as ‘driver mutations’ is dependent on the underlying mutational processes, since the probability of a particular mutation channel differs between processes. For example, a previous report has highlighted links between APOBEC-induced mutagenesis and *PIK3CA* mutations across cancer types^13^. Here we provide a comprehensive *in silico* assessment of the causal relationship between mutational processes and driver mutation acquisition across cancer types.

Selective differences between related driver mutations (e.g. differential consequences for cell evolutionary fitness) are also expected to influence the driver mutation complement of cancer samples. Traditionally it has been convenient to classify mutations found in cancer as drivers or passengers^14^, but it is likely that the effects of driver mutations actually lie on a continuum, including both ‘mini-drivers’ and major drivers^15,16^. However, the relative selective advantages of individual driver mutations have not yet been quantified. Here we analyse evidence for differential selection between frequently mutated amino acids within a driver gene by controlling for differences in the sequence-specific mutation rate, in cases where the mutational signatures alone cannot fully explain the spectra of mutations in driver genes. We also analyse differential selection between sets of related genes that show patterns of mutational exclusivity.

Together, our analysis quantifies the contributions of both mutation and selection in shaping the spectrum of driver mutations across cancer types.

## RESULTS

### Testing for mutational signature and driver mutation associations

We investigated the causal relationships between mutational signatures and mutations that caused recurrent amino acid changes within cancer types. We reasoned that when a mutational signature acts, it makes particular driver mutations, caused by certain mutation channels, more likely. We therefore tested for a difference in the levels of relative mutational signature activity in samples harbouring specific driver mutation DNA changes. The use of signature and individual channel activity information was designed to increase the sensitivity and specificity of the approach. Where the activity of a mutational signature was significantly higher in cancers with a mutation of interest compared to those without, we considered it *prima facie* evidence of a causative relationship between the mutational signature activity and the presence of the driver mutation.

Data were obtained and curated from the TCGA and International Cancer Genome Consortium (ICGC) data portals (see Methods). Driver genes were classified according to a recent study^16^. The data set for analysis represented 11,335 samples across 22 major cancer types (listed in Table S1). There were 1,448 whole genome samples and 9,887 whole exome samples. Downstream analysis was based on 14,401,296 SNVs, of which 41,201 were non-synonymous mutations in driver genes. We did not consider other types of genome alteration (such as copy number alteration).

We estimated the relative activity level (exposure) of each mutational signature in each of the samples, using non-negative least squares regression (see Methods). Simulated data showed that samples with 20 mutations or more provided sufficiently accurate recovery of mutational processes (see Methods; Figure S1). Consequently, we excluded 1,147 samples with fewer than 20 mutations from this analysis (see Methods), leaving 10,188 samples for further analysis. In each cancer type, we classed as ‘recurrent’ non-silent DNA mutations in driver genes that occurred at least four times in the cancer type. For each such change, we selected the channel among the 96 possibilities that matched the observed mutation (hereinafter the ‘causal channel’ of the change). For this channel, we identified the signatures where the frequency of the causal channel was above average, relative to all signatures active in the cancer type (Figure 1a,b,d,e). For each of these signatures, we tested for an association between signature activity and presence of the mutation in the cancer type. Power to detect associations was estimated at 14% at alpha = 0.05 (min = 0%, max = 99%), and 41/1027 tests had a power above 50% (see Methods). We found that the power was influenced by the number of times a mutation occurred, as well as the enrichment of the mutation causal channel in the signature compared to average in the cancer type (Multiple Regression, p < 2x10^-16^, 3.3x10^-9^, respectively).

**Figure 1.**
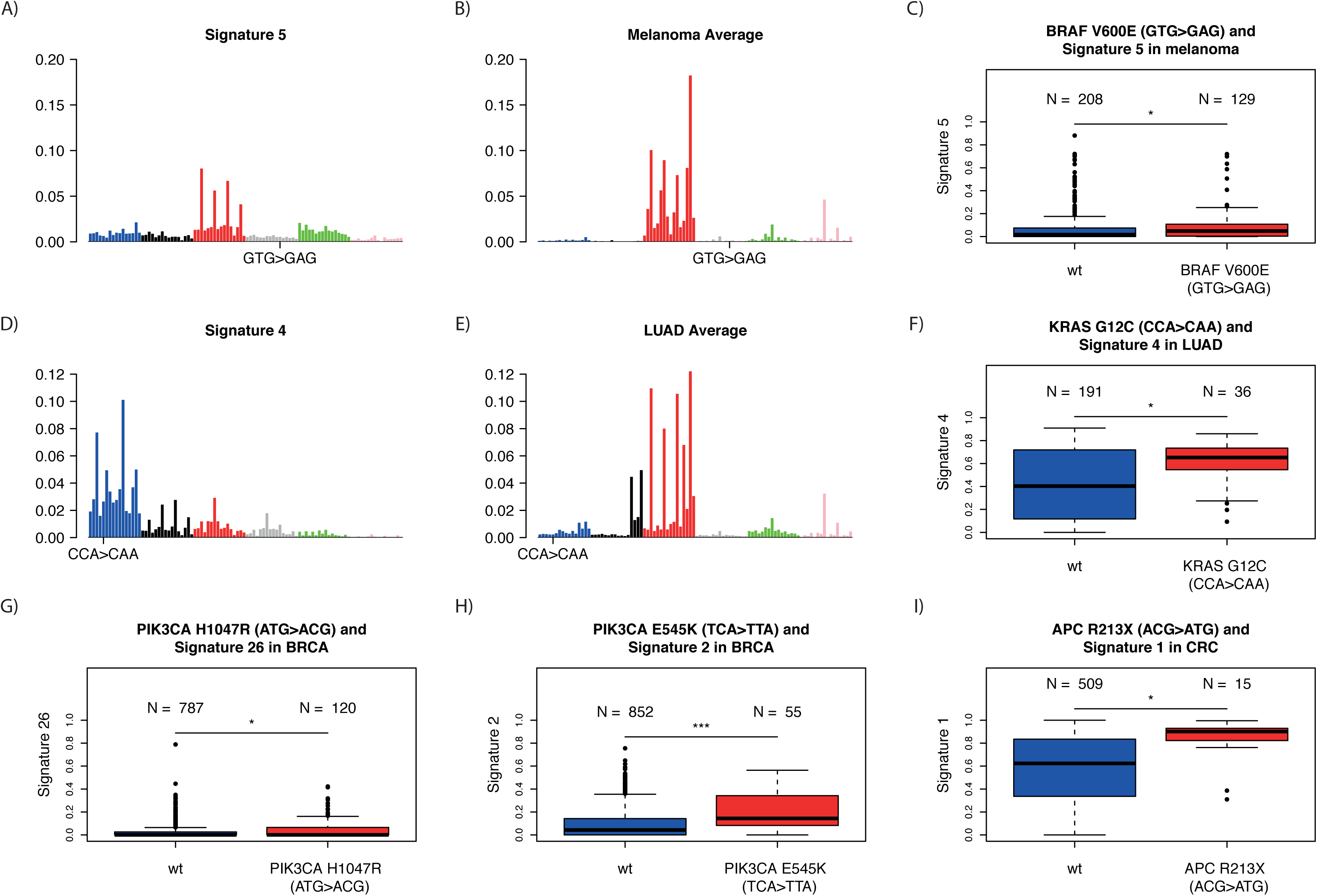
Mutational signatures active in cancer types and selected associations between mutational signature exposures and driver mutation frequencies within cancer types. **A)** Proportions of the 96 mutation channels for signature 5. **B)** Average proportions of the 96 mutation channels across all signatures reported to be active in melanoma. **C)** The signature 5 exposures are shown for melanoma samples with the *BRAF V600E* (GTG>GAG) mutation and those without. Mutated samples had significantly higher levels of signature 5. **D-F)** Same as (A-C) for signature 4 and *KRAS G12C* (ATG>ACG) in lung adenocarcinoma. **G-H)** Selected correlations between driver mutations and mutational signature exposures within cancer types. BRCA – breast cancer, CRC – colorectum, LUAD – lung adenocarcinoma.

### Mutational signatures shape driver mutation landscape

There were 56 significant associations between signature activity and driver mutations (Mann-Whitney *U* test, FDR = 0.05; one-sided test), out of 1,027 triplets of specific mutations in individual driver genes, active signatures and cancer types tested. Three of the associations involved signatures linked to extrinsic mutational processes (*i.e*. mutagens), 44 involved signatures linked to intrinsic mutational processes and 9 involved signatures with no known aetiology (Table S2 for the full list of associations).

Of the associations involving signatures linked to extrinsic mutational processes, signature 4, linked to smoking, was associated with *KRAS G12C* (CCA>CAA) in lung adenocarcinoma and with *CTNNB1* D32Y (TCC>TAC) in liver cancer (Figure 1f). Signature 24, linked to aflotoxin, was associated with *TP53 R249S* (GCC>GAC) mutations in liver cancer.

There were multiple associations involving signatures linked to intrinsic mutational processes. APOBEC activity (Signatures 2 and 13) had 17 associations. Remarkably, *PIK3CA E542K* (TCA>TTA) and *E545K* (TCA>TTA) were associated with these signatures across 7 cancer types, accounting for 76% (13/17) of the associations (Figure 1h). Additionally, *PIK3CA E453K* (TCT>TTT), and *PIK3CA E726K* (TCA>TTA) were associated with APOBEC signatures in breast cancer.

DNA mismatch repair (MMR)-linked signatures (signatures 6,15, 20 and 26) showed 15 positive associations across six cancer types (stomach, colorectum, breast, uterine carcinoma, glioma low grade and pancreas). Of these associations, *PIK3CA H1047R* (ATG>ACG) occurred three times (Figure 1g). *FBXW7 R465C* (GCG>GTG), was associated with MMR signatures in both colorectum and stomach cancer. *KRAS G12D* (ACC>ATC) and *KRAS G13D* (GCC>GTC) were associated with MMR signatures in uterine carcinoma and stomach cancer respectively. These results suggest an important role for MMR defects shaping the driver mutation spectrum of common cancers, and illustrate the likely sequence of events (early MMR-linked signatures relative to driver mutations) in some cancers with these defects.

Eleven associations with deficiency in DNA-proofreading (signature 10) were seen in uterine carcinoma and colorectum. *ARID1A R1 989X* (TCG>TTG) and *PTEN R130Q* (TCG>TTG) were each associated with this signature in both colorectum and uterine carcinoma. 3/11 positive associations involved stop-gain mutations in the *APC* gene, two in colorectum and one in uterine carcinoma (Figure 1d). Therefore it appears that (POLE) defects can cause characteristic driver lesions in these cancer types.

Seven of the associations involved signatures linked to ageing. Of particular note, signature 1 was associated with *APC R213X* (ACG>ATG) in colorectum (Figure 1i) and signature 5 was associated with *BRAF V600E* (GTG>GAG) in melanoma and *PIK3CA H1047R* (ATG>ACG) in breast cancer (Figure 1c). These results highlight the important role of ageing-related processes in cancer development.

### Detecting differential selection

The cancer genome is shaped by selection as well as mutation. We therefore tested whether selective differences between related driver mutations were discernible. To do this, we normalised for the confounding effect of mutational processes, which make specific mutations more likely than others. To implement this analysis, we grouped the samples in each cancer type into clusters of homogeneous signature exposures (Figures 2a,c,e, S2-10). After this normalisation, it is likely that within-cluster differences in driver mutation frequencies are due to selection. To test for differential selection between two related mutations in a signature-exposure cluster, we calculated the frequency of each mutation and their relative likelihoods of occurrence, inferred from the mutational signature exposures, and assuming constant mutational process activity over time. We then used the binomial test to examine the null hypothesis that the mutation counts were explained solely by their relative likelihood of occurrence (see Methods). We explored potential differential selection among the most common driver mutations (>1% of non-synonymous mutations) in nine genes: *KRAS, BRAF, NRAS, IDH1, IDH2, TP53, PIK3CA, SMAD4* and *CTNNB1* (Table S3). We conducted pairwise tests among the mutations from each gene in each sample cluster where the mutations occurred at least 10 times. Results for the same pair of mutations were combined from different samples within a cancer type using Fisher’s Method (see Methods).

**Figure 2.**
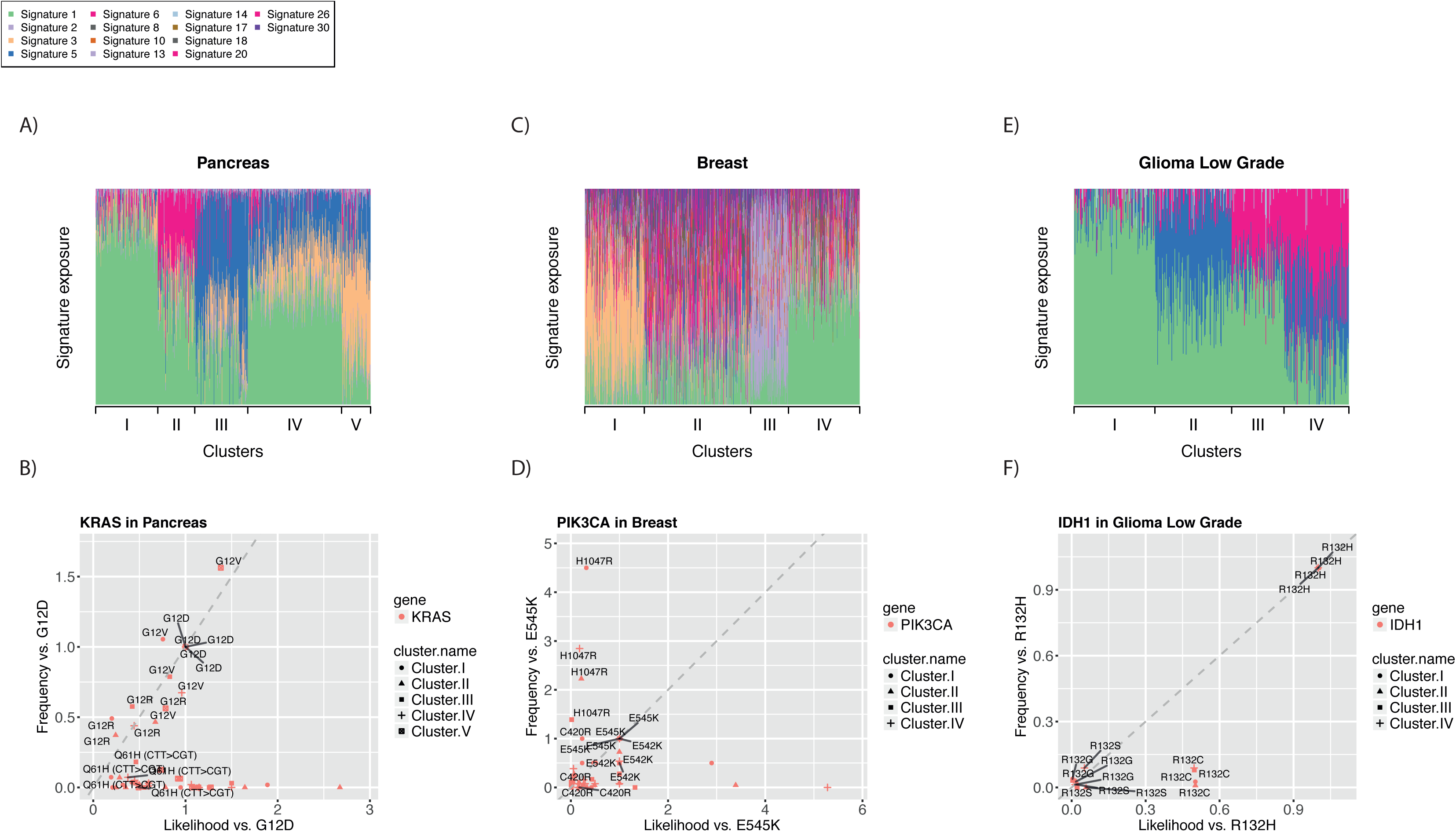
Differential selection between distinct mutations in the same driver gene. **A)** Clustering of samples based on mutational signature activities, for samples that had frequent *KRAS* mutations in pancreatic cancer. **B)** Relative observed frequency of individual *KRAS* mutations compared to *KRAS G12D* (y-axis) versus the relative mutational likelihood compared to *KRAS G12D* (x-axis), normalised by mutational cluster in pancreatic cancer (from panel A). Grey dashed line indicates expectation based on equivalent selection with *KRAS G12D* **C)** Mutational signature clustering of samples with frequent *PIK3CA* mutations in breast cancer. **D)** Relative observed frequency of *PIK3CA* mutations compared to *PIK3CA E545K* versus the relative mutational likelihood in breast cancer. **E)** Mutational signature clustering of samples with frequent *IDH1* mutations in glioma low grade. **F)** Relative observed frequency of *IDH1* mutations compared to *IDH1 R132H* versus relative mutational likelihood in glioma low grade.

### Differential selection between pathogenic amino acid changes within a driver gene

In total 10% (355/3489) of pairwise comparisons between mutations in the same gene in individual cancer types returned a significant result (Binomial Test, FDR = 0.05, Figures S2-10). In total, 8/9 genes examined had at least one pair of mutations that occurred at frequencies inconsistent with the underlying mutational likelihood (Table S4). The exception was *SMAD4*, which had the smallest number of informative mutations for this analysis.

Multiple genes and amino acid changes showed evidence of differential selection between cancer types. Among the most highly significant results, *KRAS G12D*, *G12R* and *G12V* appeared more strongly selected than other *KRAS* mutations, including *KRAS G13D*, in pancreatic cancer (Figure 2b), as did *BRAF V600E* compared to other *BRAF* common mutations, including *BRAF K601E*, in thyroid and pancreas. Also highly significant was apparent preferential selection for *PIK3CA H1047R* compared to multiple *PIK3CA* mutations, including *PIK3CA E545K* and *E542K*, in breast cancer (Figure 2d). These results suggest that there are strong selective differences among important driver mutations in the same gene in these cancer types.

A number of the results are of potential therapeutic interest. For example, we found evidence that *IDH1 R132H* is selected more strongly than *IDH1 R132C* in low grade glioma (Figure 2f). This is of particular interest given the potential specificity of therapeutic small molecular inhibitors that target *IDH1* and *IDH2* mutations^17^.

*KRAS G12C*, found to be causally related to smoking-associated signature 4 in lung adenocarcinoma (see above), also appears more strongly selected than other *KRAS* mutations (*G13D*, *A146T* and *G12S*) in this cancer type. Thus it appears that the high frequency of this *KRAS* mutation compared to others in lung adenocarcinoma is due to both smoking-associated mutational processes and the intrinsic selective advantage of the mutation.

Interestingly, the relative selective advantages of particular mutations were broadly consistent across cancer types. Specifically, out of 54 pairwise tests significant in more than one cancer type, there was only one case where a mutation that appeared selected more strongly than another in the same gene in one cancer type, but less strongly than it in another. This single case was *PIK3CA E545K* that appeared to be selected more strongly than *PIK3CA H1047R* in colorectal cancer, but below it in breast, stomach, and head and neck cancers. Overall, our results suggest that the mechanisms that underpin the selective advantage caused by a specific driver mutation are uniform across tissue types.

### Differential selection between mutationally exclusive driver genes

We next investigated differential selection between mutations within and between small sets of genes that typically show mutually exclusive mutation patterns. Using the same analysis methodology as for individual genes, we considered the common driver mutations in three sets of functionally-related genes: *KRAS*, *BRAF* and *NRAS*; *APC* and *CTNNB1*; and *IDH1* and *IDH2*.

There was evidence of greater selective differences between genes than between different residues within a gene. 3% (172/5995 pairwise comparisons) of tests were significant for mutations within a gene, whereas 14% (474/3200) were significant for mutations in different genes (Figures S11-13; Table S5). Furthermore, for two of the mutation sets - *KRAS*, *BRAF* and *NRAS* (Figure S14); and *APC* and *CTNNB1* (Figure S15) – there was significant heterogeneity across cancer types in terms of the number of mutations in each gene with evidence of preferential selection (selection above at least one other mutation in the set) (Fisher test, q = 9.6x10^-3^, 2.7x10^-3^, respectively), supporting a model where gene-specific effects on selection vary across cancer types.

Overall, all three gene sets showed evidence of differential selection in different cancer types. Amongst *KRAS*, *BRAF* and *NRAS* mutations, only particular *KRAS* mutations showed evidence of preferential selection over mutations in other genes in pancreatic cancer and uterine carcinoma (Figure 3a,b), whereas only particular *BRAF* and *NRAS* mutations showed evidence of preferential selection over mutations in other genes in melanoma and thyroid cancer (Figure 3 c,d). Illustrating this, BRAF V600E and NRAS Q61R appeared to be selected more strongly than KRAS G12D in melanoma and thyroid cancer, but more weakly than this mutation in pancreatic cancer. Other cancer types showed a range of patterns of differential selection for these three genes (Figures 3e,f, S11).

**Figure 3.**
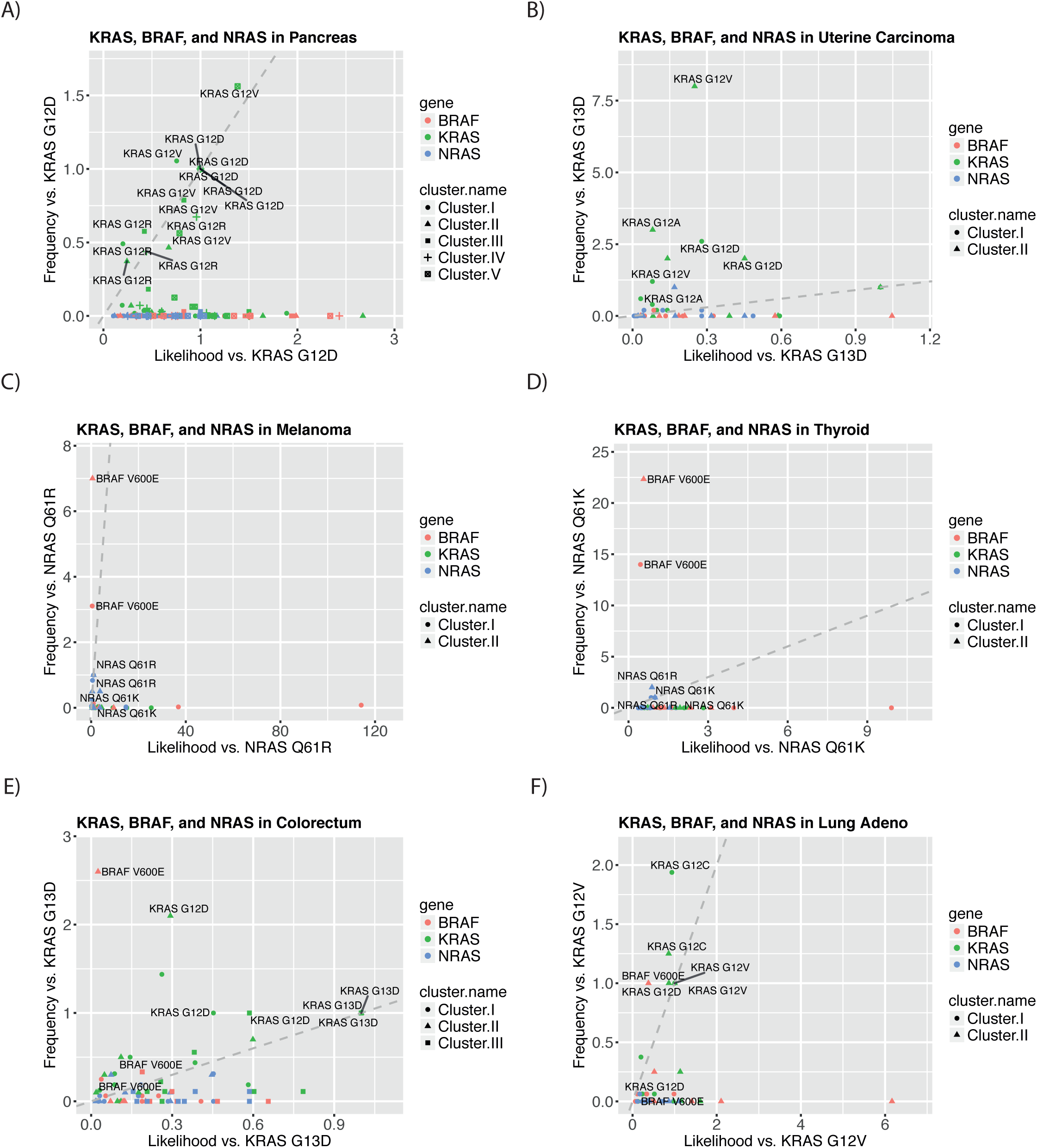
Differential selection between mutually exclusive mutations in *KRAS*, *BRAF* and *NRAS*. Scatter plots show relative observed frequencies of common driver *KRAS*, *BRAF*, and *NRAS* mutations (y-axis) plotted against relative mutational likelihood of the mutation (x-axis), where both quantities are compared to the same reference mutation. **A)** Relative observed frequencies of *KRAS*, *BRAF* and *NRAS* mutations compared to *KRAS G12D* versus relative mutational likelihoods compared to *KRAS G12D* in mutational clusters in pancreatic cancer. Grey dashed line indicates expectation based on equivalent selection with *KRAS G12D* **B)** As above, with comparison to *KRAS G13D* mutations in uterine carcinoma. **C)** As above, with comparison to *NRAS Q61R* mutations in melanoma. **D)** As above, with comparison to *NRAS Q61K* mutations in thyroid cancer. **E)** As above, with comparison to *KRAS G13D* mutations in colorectal cancer. **F)** As above, with comparison to *KRAS G12V* mutations in lung adenocarcinoma. Many mutations occur more frequently than their mutational likelihood, indicating differential (positive) selection for these mutations.

When *APC* and *CTNNB1* mutations were compared, there was evidence for selection of *CTNNB1* mutations over common *APC* mutations in each of liver cancer, uterine carcinoma, and colorectal cancer (Figure 4a,b,c). Interestingly however, evidence for selection of *APC* mutations above *CTNNB1* mutations was found in colorectal cancer only (Figure 4c).

**Figure 4.**
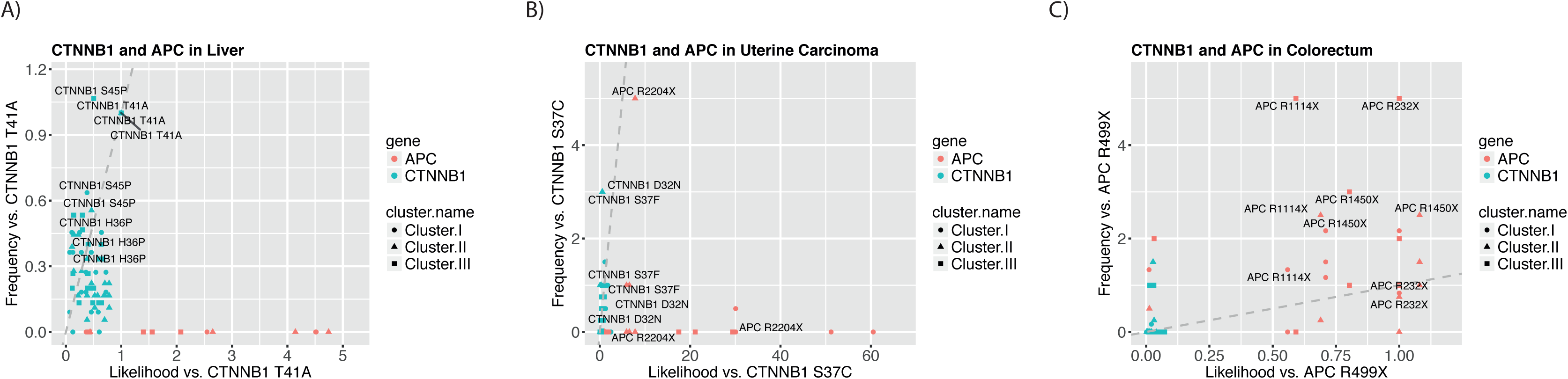
Differential selection between mutations in *CTNNB1* and *APC*. Scatter plots show relative observed mutation frequencies of *CTNNB1* and *APC* mutations (y-axis) plotted against relative mutational likelihood (x-axis), with relative frequency and likelihood by comparison to the same reference mutation. **A)** Relative frequencies of *CTNNB1* and *APC*, mutations compared to *CTNNB1 T41A* versus relative mutational likelihood compared to *CTNNB1 T41A* in sample clusters in liver cancer. Grey dashed line indicates expectation based on equivalent selection with *CTNNB1 T41A* **B)** As above, with comparison to *CTNNB1 S37C* mutations in uterine carcinoma. **C)** As above, with comparison to *APC R499X* mutations in colorectum.

Among *IDH1* and *IDH2* mutations, we found preferential selection for *IDH1 R132G* (glioma low grade) *IDH1 R132H* (glioma low grade and glioblastoma) and *IDH1 R132C* (AML and melanoma) above common *IDH2* mutations, as well as preferential selection for *IDH2 R172K* above *IDH1 R132C* in glioma low grade.

Taken together these results inform our understanding of the selective landscape and its similarities and differences between cancer types. These results suggest that both intragene and inter-gene effects contribute to differential selection, with inter-gene but not intra-gene effects varying across cancer types.

## DISCUSSION

Here, we have demonstrated associations between mutational processes and key driver mutations across cancer types, highlighting likely causative effects of these processes when they occur. Moreover, we have presented evidence in favour of selective differences between related key driver mutations across cancer types, which sheds light on the selective landscape constraining cancer evolution.

Previous work by McGranahan *et al*. examined the relationship between APOBEC associated mutational processes (signatures 2 and 13) and driver mutations and found that clonal non-synonymous mutations in driver genes occur in an APOBEC context in bladder cancer^13^. They also described subclonal mutations in driver genes in an APOBEC context in bladder, breast, head and neck, and lung cancers (uterine carcinoma and cervical cancer were not considered). Supporting their findings, we detected associations with APOBEC in bladder cancer and breast cancer, and to a lesser extent in head and neck, lung squamous, uterine carcinoma, cervical cancer and lung adenocarcinoma. Notably, we report novel associations between APOBEC activity and *ERBB2 S310F* mutations in bladder cancer. Our findings support the impression of a pervasive effect of APOBEC activity on driver mutation spectra in human cancers. Some associations we describe have been reported previously, notably the association between pack years of smoking and the *KRAS G12C* mutation in lung adenocarcinoma where the connection between the causal channel of this mutation (C>A in a CCA context) and the general tendency for tobacco carcinogens to cause transversions is well known^18,19^.

Many of the associations between mutational signatures and driver mutations presented here are novel to the best of our knowledge. Our analysis suggests that an ageing-associated process (signature 1) may cause initiating events in colorectal cancer because of the implied role of the ageing signature in causing *APC R213X* ‘gatekeeping’ mutation in colorectal cancer^16^ ^20^ – suggesting a sometimes critical role of ‘bad luck’ in this cancer type^21^. Similarly, a distinct ageing-associated process (Signature 5) appears capable of causing the common *BRAF V600E* mutation in melanoma.

Remarkably, 21/56 associations between mutational signatures and driver mutations involved *PIK3CA* mutations, and most of these associations involved signatures linked to APOBEC, which tends to occur later in carcinogenesis^13^. Thus, late arising APOBEC linked mutational processes can still have important influences on the driver mutation spectrum. Recent results showing that *PIK3CA* mutations are often subclonal^13^ support this interpretation.

Our results suggest that the selective landscape also strongly determines the driver mutation spectra. We found evidence for widespread differences in selective effects between mutations in the same gene and related genes, and moreover, that these differences appear to vary across cancer types. These results confirm that not all driver mutations have the same selective effects, and instead exist on a spectrum of selective potency. The exact nature of the selective differences we have identified will be an important area for future work. The differences we identified could reflect variation in the potential of the mutations in question to initiate disease, or alternately variation in the growth advantages conferred by these cells in established tumours. Interestingly, if there are differences in on-going growth advantages, then our data suggests that the forces of selection acting in tumours are often insufficient to displace sub-optimal mutations as less highly selected mutations remain detectable. For a limited number of driver genes, there is evidence to suggest that specific mutations correlate with disease outcomes^22,23^. Further work is needed to clarify to whether and to what extent the selective differences indicated here have prognostic and therapeutic implications.

In lung cancer, the *KRAS G12C* mutation provides a striking example of the potential for ‘alignment’ of mutation and selection: the likelihood of the *KRAS G12C* mutation is increased by smoking, but in addition it is also selectively advantageous above other common *KRAS* mutations in the disease. The same is also true for BRAF *V600E* mutations in melanoma, wherein an ageing-associated process increases the likelihood of the driver mutation, which is then subsequently strongly selected.

There are some caveats to this analysis. First, we have used data from a number of sources, which may vary in terms of quality, depth of coverage and the pipeline used to call mutations. Secondly, we have relied on the assignment of signatures to individual samples and we note that some samples have relatively few mutations, making this assignment less accurate. Relatedly, in some cancer types, there are other active signatures that were not considered in this study. Where other signatures are present, the regression method used here can only approximate the signature contributions. Thirdly, causal links between driver mutations and mutational processes are one explanation for the associations presented here, but other explanations cannot be ruled out.

In summary, our framework quantifies the combined influence of both mutation and selection on shaping a cancer’s driver mutation complement – and importantly emphasises that neither evolutionary force alone provides a sufficient explanation of the observed mutation distribution. In colon cancer for example, *BRAF* mutations (that are relatively uncommon) are mutationally unlikely, but are strongly selected. By contrast, *KRAS* drivers (that are more common), are mutationally much more likely, but are less highly selected. Our data also offer an explanation for the high frequency of driver *APC* mutations and relative paucity of driver *CTNNB1* mutations in the colon: *APC* mutations appear strongly selected and mutationally likely, whereas driver *CTNNB1* mutations are both mutationally unlikely and also less strongly selected than many *APC* driver changes. In brain cancers, the high frequency of *IDH1* mutations can be explained by strong selection for mutationally-likely *IDH1* mutations, compared to similarly mutationally likely IDH2 mutations. Overall, our results begin to delineate the distinct contributions of mutation and selection is shaping the cancer genome.

## Acknowledgements

DT acknowledges funding from the EPSRC via CoMPLEX. TG was funded by Cancer Research UK. BS-B received funding from Ludwig Cancer Research.

## METHODS

### Data collection

Mutation data (single nucleotide variants-SNVs) were downloaded from the ICGC and TCGA data portals in May 2016. We excluded data sets aligned to a reference genome other than hg19, and those with non-conforming formatting.

### Sample-specific mutation collection

Only mutations on canonical nuclear chromosomes were considered. For ICGC data, mutations labeled as ‘single base substitution’ in the simple somatic mutation files were considered for further analysis. For TCGA data, only mutations labeled as ‘SNP’ in the mutation annotation files were considered.

From these lists, non-synonymous mutations in driver genes were extracted. Driver genes definitions were as is stated below. After filtering for drivers, these mutations were re-annotated using Annovar^24^. We included mutations labeled as ‘non-synonymous SNV’, ‘stopgain’, or ‘stoploss’ in a driver gene in the annotation by Annovar.

### Definition of driver genes

Driver genes were defined using a recent study by Vogelstein et al.^16^ The list of genes is given in Table S6.

### Signature exposures for each sample in each cancer type

The 96-channel context of each SNV was imputed using the R package ‘SomaticSigantures’^25^, and the total number of SNV’s in each of the 96 channels was calculated for each sample. Mutational signatures were obtained from the Wellcome Trust Sanger Institute (http://cancer.sanger.ac.uk/cosmic/signatures) in April 2016. For whole exome data signatures were re-scaled to the trinucleotide frequencies of the exome. Non-negative least squares regression, implemented in the R package ‘nnls’^26^, was used to assign an activity score to each sample for each signature active in the applicable cancer type. Signatures activities were normalised to one for each sample to calculate the signature exposures.

### Required mutations for signature assignment

By treating each of the 30 signatures as a multinomial probability distribution, we simulated data sets from each signature with n total informative mutations (1 < n < 96). For each signature, for each value of n, we applied non-linear least squares regression to the simulated data to assign weights to the true generating signature and a set of 14 randomly chosen other signatures. We classified the regression as successful when over 50% of the regression weights were assigned to the true signature. We chose to use 15 possible generating signatures as this was above the maximum number of signatures identified in any individual cancer type. For each signature, for each number n of informative mutations, we calculated the proportion of simulations data sets where the regression was successful. We found that 20 mutations gave an average classification accuracy of 80% across signatures.

### Power calculations

We sought to test the power detect an association between mutation M, and the signature A in cancer type C, where M occurred m times in C. We considered a simple model of cancer initiation, where M is one of a set of mutations *R* of size | *R* | =n, one of which is required for cancer initiation. For these purposes we assumed n = 10.

For each random iteration of the power model we randomly selected causal channels out of 96 possibilities of the 9 other mutations in *R*. We identified the signature exposures of each sample in C. By treating the signatures as multinomial probability distributions, we then calculated the per-sample probabilities that mutation M occured rather than any of the 9 mutations in each sample. Based on these probabilities we randomly selected m samples to bear the mutation M. We then applied the Mann-Whitney U test described above.

The power was calculated as the proportion of iterations where the p-value in the test was less than the quoted value of alpha.

Out of 1,038 triplets tested where a signature represented a fold increase in the causal channel of a recurrent driver mutation in a cancer type, relatively few significant associations (56) were found. The low number of associations can be partly explained by the low average power. Even if associations were genuinely present in every case, the expected number of significant tests was 147 based on the estimated power. Indeed the significant tests were enriched for the tests with higher power (P = 3.0E-9, Mann-Whitney *U* test, mean power among significant tests and non-significant tests, 32% and 13%, respectively). Part of the reason for this is the technical challenges inherent in deconvolving mutational signature intensities. Timing mismatches between the activity of a mutational selection and the window of selection for a driver mutation probably also contribute to the low numbers of associations.

### Clustering of samples into groups of homogeneous signature exposure

Samples within a cancer type were clustered based on their signature exposures using the R package ‘fpc’^27^. The optimum number of clusters, between one and the number of signatures reported in the cancer type, was selected using the average silhouette width criterion.

### Testing for evidence of differential selection between mutations in a cluster of samples

For a sample cluster S, we sought to test the null hypothesis that the frequencies, m_1_ and m_2_, of two mutations, M_1_ and M_2_, were consistent with their likelihoods of occurrence. We considered a meta-sample with an exposure for each signature equal to the average exposure for that signature across the samples in S. Based on the signature exposures of the meta-sample, treating each signature as a multinomial probability distribution, we calculated, l_1_ and l_2_, the probability of the causal channels of M_1_ and M_2_ respectively occuring among the 96 mutation types.

We used a binomial test to test whether the relative frequencies of M_1_ and M_2_ were consistent with their relative likelihoods of occurrence. Specifically we found the probability, p_greater_ that X > m_1_, and the probability p_less_ that X < m_1_ where:

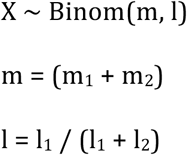

### Testing for evidence of differential selection between mutations in a cancer type

For mutations M_1_ and M_2_ in sample clusters Si (I in 1,…n) in cancer type C, we found the probabilities p_i,greater_ and p_i,less_, for each cluster, as described above. The separate values of p_i,greater_ were combined for the different sample clusters using Fisher’s Method to give p_greater_, as were the separate values of p_i,less_ to give p_less_. P_greater_ and p_less_ were combined to give a single pvalue, p, for the comparison using the formula

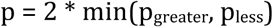

